# The Histone Methyltransferase KMT2D promotes Natural Killer cell effector molecule release

**DOI:** 10.64898/2026.09.16.752157

**Authors:** Katelynn R. Kazane, Jeong Hyun Ji, Georgia R. Lill, Sebastian Zolog, Christian G. Bustillos, David Hsieh, Evelyn Hernandez-Almonte, Luis A. Pedroza, Maureen A. Su, Jordan S. Orange, Timothy E. O’Sullivan

## Abstract

Natural Killer (NK) cells eliminate virally infected and cancerous cells by secreting cytotoxic granules and pro-inflammatory cytokines. However, the epigenetic mechanisms that coordinate these processes in human NK cells remain poorly understood. We identified two NK deficiency (NKD) patients with mutations in the histone methyltransferase KMT2D that demonstrated impaired cytotoxicity and degranulation. CRISPR-mediated KMT2D deletion or introduction of a patient-specific mutation in healthy human NK cells was sufficient to inhibit effector functions. NK cell-specific KMT2D deletion in mice resulted in defective NK cell degranulation and IFN-γ production, increasing mortality following MCMV infection. KMT2D loss reduced H3K4me1 deposition, associated with decreased levels of RAB3D. RAB3D-deficient human NK cells reduced the release of GZMB and IFN-γ, without impacting intracellular levels. Thus, KMT2D acts as a conserved epigenetic regulator of mature NK cell functions, promoting rapid effector molecule release.

**HIGHLIGHTS:**

- Mutations in KMT2D are associated with human NKD.
- KMT2D is required for NK cell cytotoxicity and degranulation.
- Loss of KMT2D in mouse NK cells increases susceptibility to MCMV infection.
- KMT2D regulates RAB3D to enhance the release of effector molecules.

## INTRODUCTION

Natural Killer (NK) cells protect against viral infections and cancer^1,2^ through two distinct effector pathways: the production of pro-inflammatory cytokines such as Interferon (IFN)-γ, and the polarized release of perforin and granzyme-containing lytic granules^3,4^. While both effector pathways require vesicle trafficking and exocytosis to release their contents, each uses distinct spatial routes^5^. Lytic granules are directed toward the immune synapse following microtubule-organizing center polarization, whereas cytokines are released in multiple directions via pathways involving the recycling endosome^5,6^. Despite the importance of both processes for NK cell immunity, whether these are regulated by common transcriptional or epigenetic programs remains poorly understood.

Inborn errors of immunity in NK cell effector function or development are associated with human NK cell deficiencies (NKDs) characterized by recurrent herpesvirus infections^7,8^. Genetic analyses of NKDs and other combined immunodeficiencies have identified genes encoding transcription factors, cell cycle regulators, and components of cytotoxic granule trafficking, release, and biogenesis such as PRF1, RAB27A, UNC13D, and STXBP2^9,10^. However, candidate genes nominated from the Natural Killer Cell Evaluation and Research (NEAR) cohort following clinical phenotyping and whole-exome sequencing have no known role in human NK cell function^10^. Analysis of the NEAR cohort candidate genes identified KMT2D, a COMPASS-family lysine methyltransferase that regulates enhancer activity and is associated with monomethylation of histone H3 lysine 4 (H3K4me1)^11,12^, suggesting that NKDs may also arise from germline mutations in epigenetic regulators.

Here, we demonstrate that KMT2D is a conserved epigenetic regulator of NK cell cytotoxicity and degranulation in mice and humans, in addition to mouse NK cell antiviral function. Loss of KMT2D reduced H3K4me1 levels in human NK cells and resulted in decreased expression of a conserved set of genes associated with exocytosis and vesicle-mediated trafficking in mouse and human NK cells. Specifically, KMT2D enhanced levels of the vesicle-associated GTPase RAB3D, which was required for optimal lytic granule degranulation and cytokine release. Together, these findings suggest that KMT2D-mediated regulation of RAB3D supports the two main secretory pathways involved in NK cell effector function, and that loss of KMT2D in NK cells increases susceptibility to viral infection in mice.

## RESULTS

### KMT2D is required for NK cell cytotoxicity

To identify epigenetic regulators associated with human NKD, we utilized whole-exome sequencing data from 99 patients in the published NEAR cohort^10^. Of these patients, 29 individuals displayed variants linked to known regulators of NK cell function (solved), 17 displayed candidate variants in genes not previously implicated in NK function (candidate), and 53 had no identifiable genetic component (unsolved) (**Figure 1A**). Candidate genes included 3 transcription factors, 2 transcriptional repressors, and KMT2D as the sole epigenetic regulator. Peripheral blood mononuclear cells (PBMCs) from two KMT2D variant patients displayed impaired cytotoxicity and contained lower frequencies of circulating NK cells than healthy controls (**Figures 1B,C**). Both patients carried heterozygous variants in *KMT2D*, one missense variant and one deletion (**Figures 1D,S1A**). Despite the reduced frequencies of NK cells, analysis of NK cell maturation by flow cytometry revealed only a modest decrease in the frequency of terminally differentiated CD16^+^CD57^+^ NK cells in one patient compared to a maternal control (**Figures S1B,C**). While these observations implicated KMT2D variants in altered human NK cell development and impaired cytotoxicity, whether KMT2D is required for cytotoxicity in mature human NK cells was unknown. To test the cell-intrinsic requirement for KMT2D in mature NK cell cytotoxicity, we isolated mature NK cells from healthy donors and generated *KMT2D* knockout (*KMT2D* KO) cells and non-targeting controls (NT) using Cas9 adenine base editing^13^ (**Figures S1D-F**). Following co-culture with K562-GFP^+^ leukemia cells, *KMT2D* KO human NK cells displayed significantly less killing than control-edited NK cells (**Figure 1E**). Furthermore, *Kmt2d* Cas9-ribonucleoprotein (cRNP) edited mouse NK cells (*Kmt2d* KO) (**Figures S1G,H**) showed a significant reduction in killing of B2M-deficient MC38 tumor cells compared to *Rosa26*-edited controls (**Figure 1F)**. These results demonstrate a conserved, cell-intrinsic requirement for KMT2D in NK cell cytotoxicity, independent of NK cell development.

**Figure 1.**
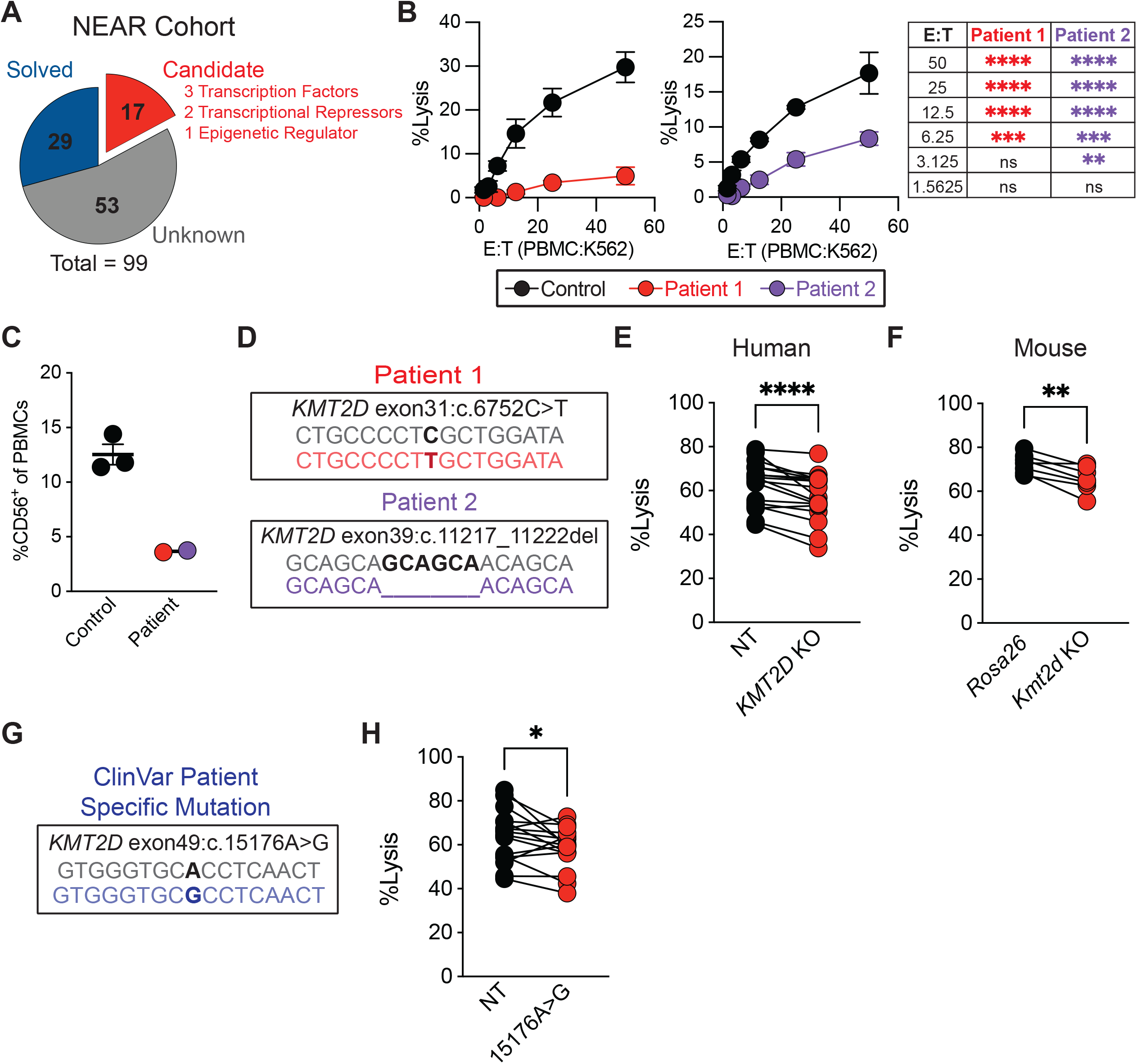
KMT2D is required for human NK cell cytotoxicity. (**A**) Pie chart depicting the proportion of NEAR participants with solved, candidate, or unsolved genetic causes of NKD as reported in Abdalgani et al.^10^. (**B**) Percent specific lysis of K562 tumor cells after co-culture with patient or unaffected donor (Control) PBMCs at varying Effector:Target (E:T) ratios. (**C**) Percent of CD56^+^ cells from patient versus Control PBMCs. (**D**) Sequence alignment between KMT2D-deficient patients and GRCh38 assembly. (**E**) Percent specific lysis of K562-GFP tumor cells after 16hr co-culture with Cas9 base edited human NK cells with either a non-targeting (NT) or *KMT2D*-specific (*KMT2D* KO) guide RNA at a 1:4 E:T in the presence of IL-2 and IL-15. (**F**) Percent specific lysis of B2M-deficient MC38 tumor cells after 16hr co-culture with cRNP edited mouse NK cells with guide RNA targeting either *Rosa26* or *Kmt2d* (*Kmt2d* KO) at a 1:2 E:T in the presence of IL-15. (**G**) Sequence alignment between ClinVar identified likely pathogenic mutation c.15176A>G and GRCh38 assembly. (**H**) Percent specific lysis of K562-GFP tumor cells after 16hr co-culture with Cas9 base edited human NK cells with 15176A>G mutation versus NT at a 1:4 E:T in the presence of IL-2 and IL-15. Data represent (**B**) mean ± SEM of three technical replicates from a single donor, (**C**) individual donors, or (**E,F,H**) individual paired donors or mice and at least two independent experiments with (**E**) n = 17 or (**H**) n = 16 human donors, or (**F**) n = 7 mice. *p<0.05, **<0.01, ***<0.001, ****<0.0001 by (**B**) two-way ANOVA or (**E,F,H**) Wilcoxon matched-pairs sign rank test.

Given that two germline variants in KMT2D are associated with human NKD, we tested whether an independently reported likely pathogenic missense KMT2D variant from the NIH ClinVar database^14^ was sufficient to impair human NK cell cytotoxicity (**Figure 1G**). Introduction of this variant into healthy donor mature human NK cells significantly impaired target-cell killing relative to non-targeting controls (**Figure 1H**). Together, these results associate heterozygous KMT2D variants with NKD and demonstrate that loss of KMT2D impairs mammalian NK cell cytotoxicity.

### KMT2D is required for mouse NK cell antiviral responses

Since recurrent severe human cytomegalovirus infections are a hallmark of NKD^7^, we interrogated the role of KMT2D in mouse NK cells during mouse cytomegalovirus (MCMV) infection. NK cell-specific deletion of *Kmt2d* (*Ncr1*^Cre/+^*Kmt2d*^fl/fl^; NK*-Kmt2d* KO) in mice resulted in a greater loss of body weight starting at day 2 post-infection (p.i.) compared to littermate controls (*Ncr1*^+/+^*Kmt2d*^fl/fl^; WT), resulting in a significant survival deficit in NK-*Kmt2d* KO mice by day 7 p.i. (**Figures 2A-C**). Given these results, we then tested whether heterozygous loss of *Kmt2d* in mouse NK cells resulted in a cell-intrinsic decrease in effector function during viral infection. We generated mixed bone marrow chimeric mice (mBMC) with bone marrow from NK-specific heterozygous *Kmt2d*-deficient mice (*Ncr1*^Cre/+^*Kmt2d*^fl/+^ CD45.2; *Kmt2d*^+/-^) and WT (CD45.1) mice, adoptively transferred into WT CD45.1 x CD45.2 hosts, and infected them with MCMV (**Figure 2D**). Across all tissues analyzed on day 1.5 p.i., there was a significant decrease in the frequency and amount of IFN-γ and perforin-producing NK cells in *Kmt2d*^+/-^compared to WT controls, respectively (**Figures 2E,F**). These results were not due to developmental defects in mouse NK cells, as we observed a significant increase in both the proportion and number of circulating NK cells from the blood of NK-*Kmt2d* KO mice (**Figures S2A,B)**. Furthermore, there was a significant increase in the number of mature (KLRG1^+^ CD27^-^NK1.1^+^) and Ly49H^+^ NK cells in the blood of NK-*Kmt2d* KO versus WT mice (**Figures S2C,D**), with similar results found in *Kmt2d*^+/-^NK cells analyzed from the blood and peripheral tissues of WT:*Kmt2d*^+/-^mBMC mice **(Figures S2E-H)**. Thus, while *Kmt2d* limits the abundance of mature mouse NK cells during development, the defects in effector molecules and enhanced viral susceptibility observed with NK cell-specific *Kmt2d* loss were likely due to defects in mature NK cell production of effector molecules.

**Figure 2.**
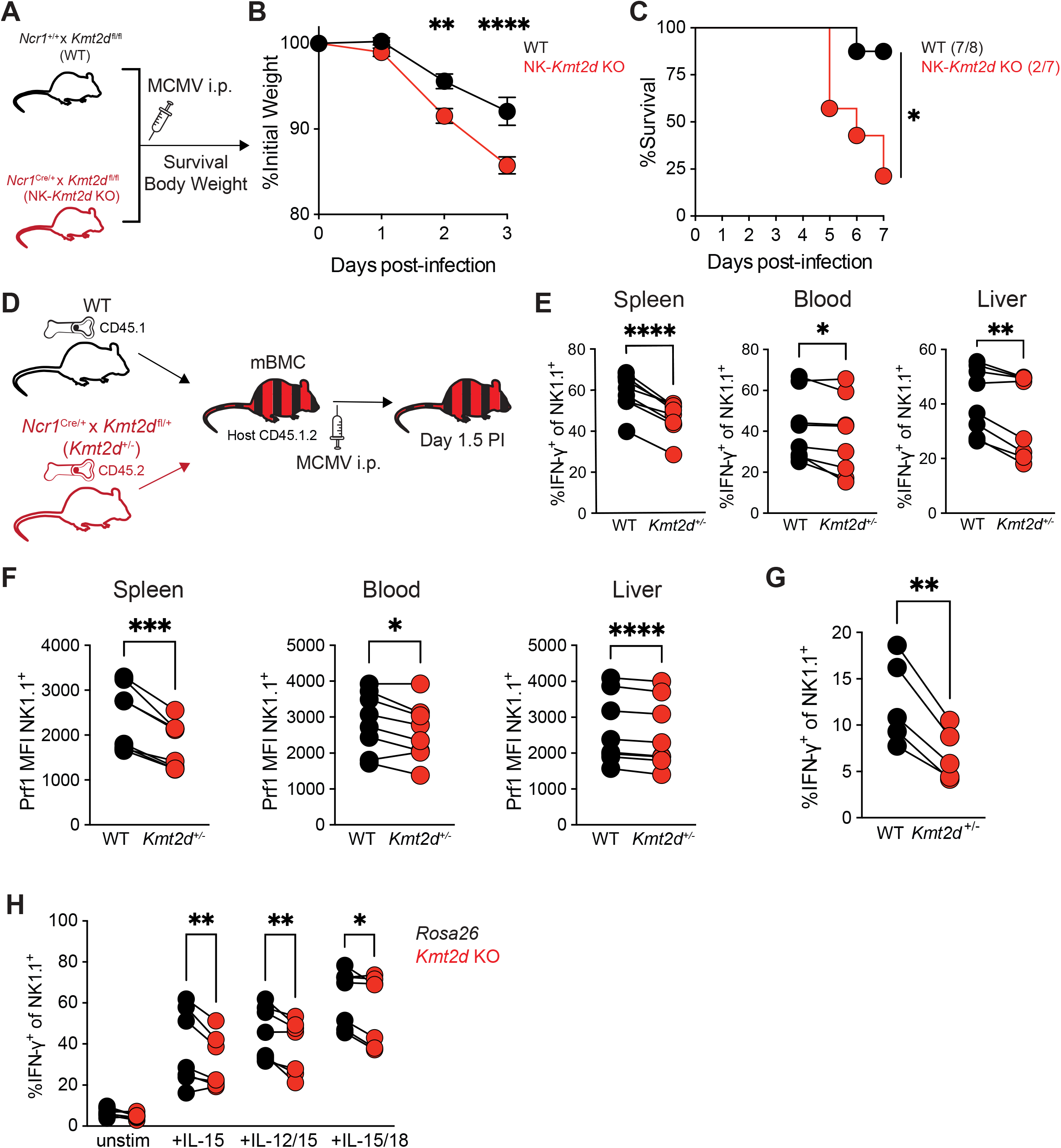
*Kmt2d* is required for NK antiviral responses during MCMV infection. (**A**) Diagram of *Ncr1*^+/+^*Kmt2d*^fl/fl^ (WT) versus *Ncr1*^Cre/+^*Kmt2d*^fl/fl^ (NK-*Kmt2d* KO) MCMV infection experiments. (**B,C**) Percent of initial body weight (**B**) and survival (**C**) of WT versus NK-*Kmt2d* KO mice challenged with MCMV at specified days post-infection (p.i.). (**D**) Schematic of mixed-bone marrow chimeric (mBMC) mouse generation from WT and *Ncr1*^Cre/+^*Kmt2d*^fl/+^ (*Kmt2d^+/-^)* mice followed by MCMV challenge. (**E,F**) Percent IFN-γ^+^ (**E**) and mean fluorescence intensity (MFI) of perforin (Prf1) (**F**) of NK1.1^+^ WT versus *Kmt2d*^+/-^cells isolated from the spleen (left), blood (middle), and liver (right) of mBMCs at day 1.5 p.i.. (**G**) Percent IFN-γ^+^ of NK1.1^+^ WT versus *Kmt2d*^+/-^cells isolated from uninfected mBMCs after anti-NK1.1 stimulation. (**H**) Percent IFN-γ^+^ of NK1.1^+^ *Rosa26* versus *Kmt2d* KO mouse NK cells after 16 hr incubation with indicated cytokines. Data represent (**B,C**) mean ± SEM or (**E,H**) paired individuals, and at least two independent experiments with (**B**) n=9, (**C**) n=7-8, (**E,F**) n=8, (**G**) n=5, or (**H**) n=7 mice. *p<0.05, **<0.01, ***<0.001, ****<0.0001 by (**B**) two-way ANOVA, (**C**) Mantel-Cox test, (**E,G**) Wilcoxon matched-pairs sign rank test, or (**H**) paired two-way ANOVA.

To test this hypothesis, we isolated splenic NK cells from uninfected WT:*Kmt2d*^+/-^mBMC mice and activated them with plate-bound anti-NK1.1 antibodies. We observed a significant decrease in the frequency of IFN-γ in *Kmt2d*^+/-^NK cells following stimulation (**Figure 2G**). We then isolated mature NK cells from the spleen of uninfected WT mice and generated *Kmt2d* KO or control *Rosa26* edits using cRNPs. Following pro-inflammatory cytokine stimulation, the proportion of IFN-γ^+^ NK cells was significantly lower in *Kmt2d* KO versus *Rosa26*-edited NK cells (**Figure 2H**). Altogether, these data suggest that *Kmt2d* regulates the antiviral function of mature mouse NK cells during MCMV infection.

### KMT2D maintains an H3K4me1-associated transcriptional program in human NK cells

To define the conserved transcriptional programs regulated by KMT2D in NK cells independent of development, we performed RNA sequencing (RNA-seq) on splenic NK cells isolated from NK-*Kmt2d* KO and WT mice and 10 (5 male and 5 female) healthy donor-matched control-edited and *KMT2D* KO NK cells (**Figures S3A,B**). While NK-*Kmt2d* KO mouse NK cells exhibited broad transcriptional changes, with 868 genes downregulated and 688 genes upregulated relative to WT NK cells (**Figure 3A**), donor-matched *KMT2D* KO NK cells displayed 242 downregulated and 115 upregulated genes (**Figure 3B**). Comparison of the differentially expressed genes in mouse and human KMT2D-deficient NK cells identified 52 shared genes (**Figure 3C**), suggesting either significant species-specific differences or an enrichment of KMT2D-regulated genes in mature NK cells. Functional enrichment analysis of the conserved downregulated genes identified three terms related to effector molecule release: secretory granule, exocytosis, and regulation of vesicle-mediated transport (**Figure 3D**). However, transcripts encoding the effector molecules IFN-γ, perforin (PRF1), or granzyme B (GZMB) were not reduced in the absence of KMT2D **(Figures S3C-E)**. Therefore, these results suggested that the functional defects observed in KMT2D-deficient NK cells could be associated with decreased expression of genes involved in effector molecule release.

**Figure 3.**
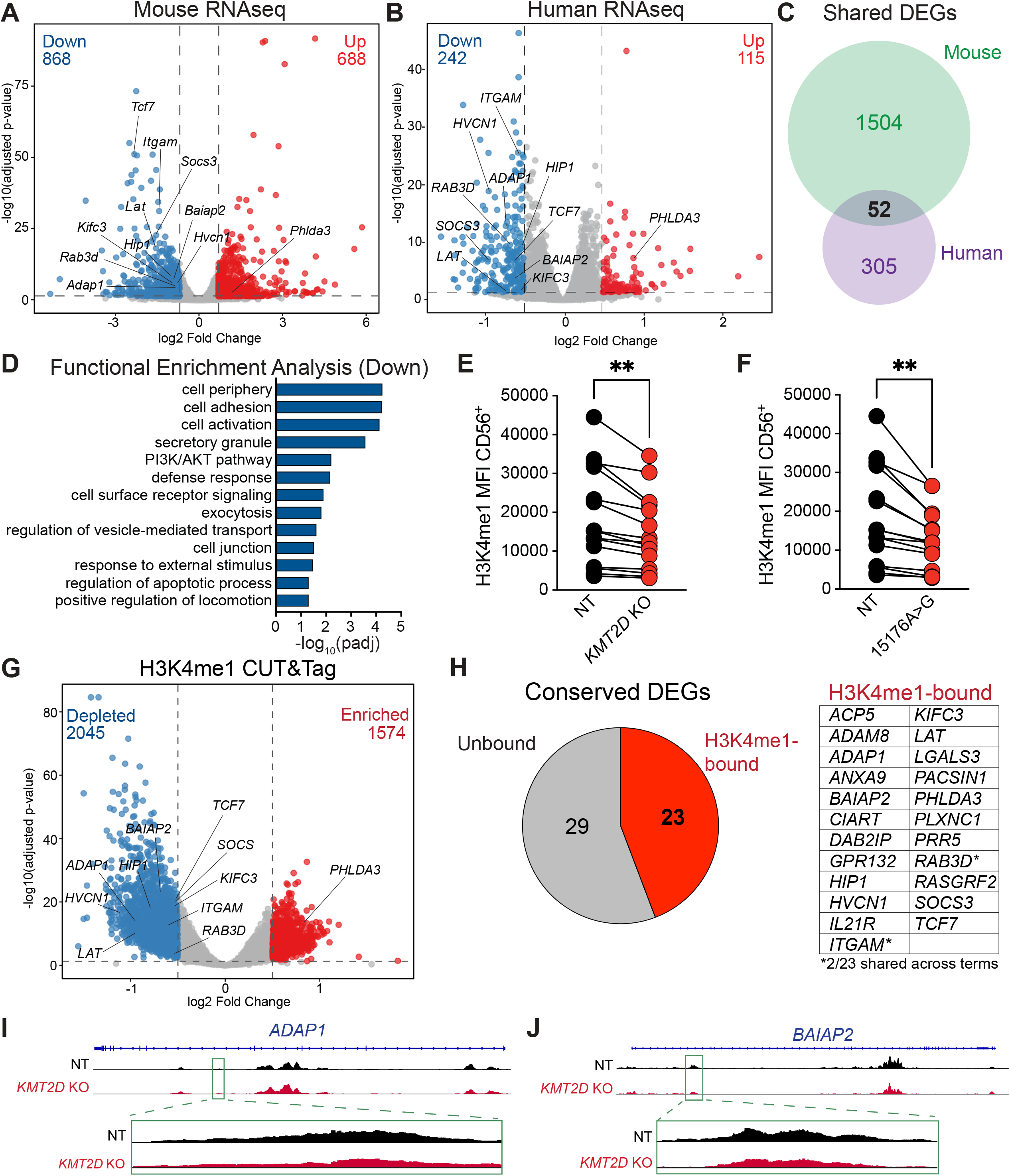
KMT2D regulates NK cells via H3K4me1 deposition. (**A,B**) Volcano plots showing significant differentially expressed genes (DEGs) from RNA sequencing (RNA-seq) of *Kmt2d* KO versus WT mouse NK cells (**A**), or *KMT2D* KO versus NT edited human NK cells (**B**). (**C**) Venn diagram showing the overlap of directionally conserved DEGs in mouse and human RNA-seq. (**D**) Functional enrichment analysis of overlapping mouse and human DEGs. (**E,F**) H3K4me1 MFI in CD56^+^ NT versus *KMT2D* KO (**E**) or 15176A>G (**F**) human NK cells. (**G**) Volcano plot showing significant differentially bound peaks from H3K4me1 CUT&Tag of *KMT2D* KO versus NT human NK cells. (**H**) Chart showing the number of H3K4me1-bound versus unbound overlapping RNA-seq DEGs (left) and the list of bound genes (right). (**I,J**) Representative tracks showing anti-H3K4me1 CUT&Tag peaks at the *ADAP1* (**I**) and *BAIAP2* (**J**) loci in *KMT2D* KO versus NT human NK cells. Data represent (**A,B**) individual genes from RNA-seq (**G**) individual peaks from H3K4me1 CUT&Tag, or (**E,F**) individual human donors with (**A**) n=3 mice, (**B**) n=10, (**E,F**) n=15, or (**G**) n=3 donors and (**E,F**) at least two independent experiments. *p<0.05, **<0.01 by Wilcoxon matched-pairs sign rank test.

KMT2D acts as an epigenetic regulator through its methyltransferase activity facilitating H3K4me1 deposition^11,15^ and its association with the histone demethylase UTX^16^. To discern through which mechanism KMT2D regulated transcriptional programming in human NK cells, we examined whether loss of KMT2D altered global H3K4me1 abundance. *KMT2D* KO or introduction of the KMT2D c.15176A>G variant in healthy donor human NK cells reduced global H3K4me1 relative to non-targeting controls (**Figures 3E,F**). To determine whether this loss was accompanied by locus-specific changes in chromatin modifications, we performed CUT&Tag profiling^17^ of H3K4me1 and H3K27ac in donor-matched *KMT2D* KO and control-edited human NK cells. *KMT2D* KO resulted in depletion of 2,045 H3K4me1 peaks and enrichment of 1,574 peaks relative to control-edited NK cells (**Figure 3G**). Most differential H3K4me1 peaks were located within intronic regions (57.2%) or distal intergenic regions (23.8%) (**Figure S3F**). Of the 52 shared DEGs in human and mouse KMT2D-deficient NK cells, 23 were associated with differential H3K4me1 peaks in human NK cells (**Figure 3H**). Differential H3K27ac peaks overlapped more extensively with KMT2D-dependent H3K4me1 peaks than with genes differentially expressed following KMT2D loss (**Figure S3G**). These findings indicate that KMT2D-dependent changes in enhancer-associated histone modifications are not uniformly associated with altered steady-state gene expression and raise the possibility that they instead influence stimulus-dependent transcriptional responses.

Since UTX is stabilized by KMT2D^18^ and contributes to sex-dependent differences in effector-molecule production by mouse and human NK cells^19^, we tested whether conserved KMT2D-dependent genes were regulated by loss of UTX. Of the conserved KMT2D-dependent genes, 24 were reported to be differentially expressed in UTX-deficient mouse NK cells^19^, and of those, 12 were bound by UTX in mouse NK cells (**Figure S3H**). However, while 2 UTX-bound genes, *ADAP1* and *BAIAP2,* were associated with cell surface signaling or secretory granule gene ontology terms, respectively, CRISPR-mediated deletion of these genes in mature human NK cells showed no difference in tumor killing compared to non-targeting controls (**Figures S3I-L**). Furthermore, while KMT2D deletion reduced UTX levels in female, but not male, donor NK cells, KMT2D loss impaired cytotoxicity in NK cells from both female and male donors (**Figures S3M,N**). These results suggest that decreased UTX levels in *KMT2D* KO female NK cells are not sufficient to reduce cytotoxicity in the absence of KMT2D. Additionally, H3K4me1 levels were decreased in *KMT2D* KO NK cells from both female and male donors (**Figure S3O**), suggesting a sex-independent association between KMT2D function, global chromatin H3K4me1 levels, and human NK cell cytotoxicity.

### RAB3D is required for release of effector molecules in human NK cells

To identify KMT2D-regulated genes required for cytotoxicity in human NK cells, we selected genes associated with multiple secretory pathway-related terms from gene ontology analysis whose loci were differentially bound by H3K4me1. Only *RAB3D* and *ITGAM* (CD11b) were associated with secretory granule, exocytosis, and regulation of vesicle-mediated transport terms (**Figure 3D**). While CD11b promotes NK cell adhesion^20^, and we observed decreased surface expression of CD11b in *KMT2D* KO NK cells (**Figure S4A**), the proportion of NK-tumor conjugates formed by *KMT2D* KO versus NT NK cells was not impaired during killing assays (**Figure S4B**). Therefore, we prioritized the GTPase RAB3D due to its association with secretory granules in mast cells^21^ and the broader functions of RAB3-family proteins in vesicle organization and regulated exocytosis in neurons and endocrine cells^22,23^. RAB3D transcripts and protein levels were significantly reduced in *KMT2D* KO human NK cells (**Figure 4A,B**), with reduced H3K4me1 occupancy at the RAB3D locus following KMT2D deletion (**Figure 4C**). Furthermore, CRISPR-edited *RAB3D*-deficient (*RAB3D* KO) human NK cells displayed significantly reduced tumor killing compared to non-targeting controls (**Figure 4D**), suggesting that KMT2D-dependent regulation of RAB3D levels was required for optimal lytic granule degranulation and cytokine release.

**Figure 4.**
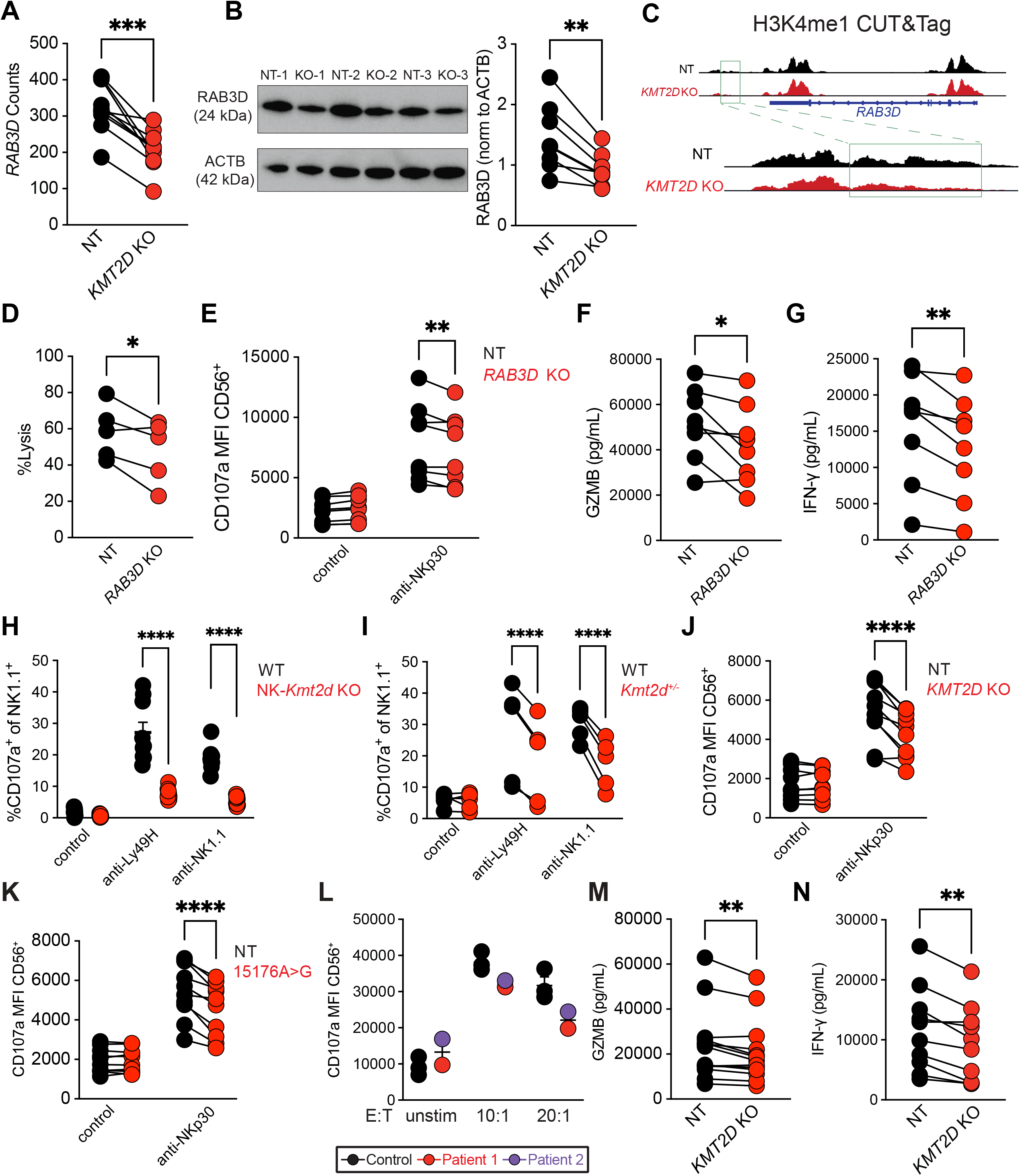
KMT2D regulates RAB3D to control NK degranulation and cytokine secretion. (**A**) *RAB3D* transcript counts in *KMT2D* KO versus NT human NK cells from RNA-seq. (**B**) Immunoblot and quantification of RAB3D normalized to ACTB in NT versus *KMT2D* KO human NK cells. (**C**) Representative tracks showing anti-H3K4me1 CUT&Tag peaks at the *RAB3D* loci in *KMT2D* KO versus NT human NK cells. (**D**) Percent specific lysis of K562-GFP tumor cells after 16hr co-culture with cRNP-edited *RAB3D* (*RAB3D* KO) versus NT human NK cells at a 1:4 E:T in the presence of IL-2 and IL-15. (**E**) CD107a MFI in CD56^+^ *RAB3D* KO versus NT human NK cells after 16 hr stimulation with plate-bound anti-NKp30 or control conditions in the presence of IL-2, IL-15 and 4 hours of anti-CD107a, Brefeldin A (BFA), and Monensin. (**F,G**) Quantification via ELISA of secreted GZMB (**F**) or IFN-γ (**G**) by NT versus *RAB3D* KO human NK cells after 16 hr co-culture with K562 tumor cells, IL-2, IL-15, and IL-18. (**H,I**) Percent CD107a^+^ of NK1.1^+^ WT versus NK-*Kmt2d* KO (**H**) or *Kmt2d*^+/-^mBMC (**I**) splenocytes after 4 hr stimulation with plate-bound anti-Ly49H, plate-bound anti-NK1.1, or in control wells co-cultured with IL-15, anti-CD107a, BFA, and Monensin. (**J-L**) MFI of CD107a in CD56^+^ *KMT2D* KO (**J**) or 15176A>G (**K**) versus NT, or Patient versus Control (**L**) human NK cells after 16 hr (**J,K**) or 4hr (**L**) stimulation with plate-bound anti-NKp30 (**J,K**), K562 tumor cells (**L**) or control conditions in the presence of IL-2, IL-15 and 4 hours of anti-CD107a, BFA, and Monensin. (**M,N**) Quantification via ELISA of secreted GZMB (**M**) or IFN-γ (**N**) by *KMT2D* KO versus NT human NK cells after 16 hr co-culture with K562 tumor cells, IL-2, IL-15, and IL-18. Data represent paired individuals **(A,B,D-G,I-K,M,N**) or individual mice (**H**) and at least two independent experiments, or independent donors (**L**) with (**A**) n=10, (**B,D-G**) n=8, (**J,K**) n=10, or (**M,N**) n=13 donors or (**H**) n=10 or (**I**) n=5 mice. *p<0.05, **<0.01, ***<0.001, ****<0.0001 by (**A,B,D,F,G,M,N**) Wilcoxon matched-pairs sign rank test, (**E,I,J,K**) paired two-way ANOVA, or (**H**) unpaired two-way ANOVA.

To test this hypothesis, we stimulated *RAB3D* KO and NT human NK cells with anti-NKp30 in the presence of anti-CD107a (LAMP-1) antibodies to quantify lytic granule fusion with the surface of NK cells via flow cytometry. While stimulated *RAB3D* KO human NK cells displayed a significant decrease in surface CD107a compared to NT controls (**Figure 4E**), no significant differences were observed in intracellular IFN-γ, PRF1, or GZMB levels (**Figures S4C-E**), suggesting a defect in effector molecule release. In support of this hypothesis, quantification of effector molecules released by NK cells via Enzyme-Linked Immunosorbent Assay (ELISA) following cytokine stimulation revealed a significant decrease in the amount of GZMB and IFN-γ in the supernatant of *RAB3D* KO versus NT human NK cells (**Figures 4F,G**). Together, these data suggest that RAB3D is required for the optimal release of effector molecules from human NK cells.

Given these results, we then investigated whether *KMT2D*-deficient NK cells similarly showed defects in the release of effector molecules. Plate-bound antibody-stimulated mouse NK-*Kmt2d* KO and *Kmt2d*^+/-^NK cells, human *KMT2D* KO, 15176A>G mutant, and both *KMT2D* variant NKD patient NK cells all displayed significantly decreased CD107a levels compared to controls (**Figures 4H-L**). Similar to *RAB3D* KO cells, we did not observe significant differences in intracellular PRF1 or GZMB levels following stimulation of either *KMT2D* KO or 15176A>G mutant NK cells (**Figures S4F-I**). However, *KMT2D* KO human NK cells released significantly less GZMB and IFN-γ into supernatants than controls (**Figures 4M,N**), suggesting that KMT2D loss results in defective release of effector molecules from human NK cells, rather than reducing the levels of effector molecules. Thus, these data demonstrate that KMT2D-dependent epigenetic regulation of RAB3D enhances human NK cell effector functions through control of effector molecule release, with implications for human inborn errors of immunity.

## DISCUSSION

Our study identifies KMT2D as an epigenetic regulator of NK cell effector functions and implicates KMT2D in immunodeficiencies with loss of NK cell function. Two individuals in the NEAR cohort carrying heterozygous KMT2D variants exhibited reduced NK cell abundance, impaired PBMC-mediated cytotoxicity, and defective NK cell degranulation. KMT2D deletion in mature human and mouse NK cells resulted in similar decreases in degranulation, cytokine production, and cytotoxicity, establishing a conserved, cell-intrinsic requirement for KMT2D in NK cell effector functions independent of developmental effects. Furthermore, NK cell-specific *Kmt2d* deficiency impaired effector responses during MCMV infection and increased infection-associated weight loss and mortality. These findings expand the genetic framework of human immune deficiencies beyond mutations in proximal components of the cytolytic machinery, such as PRF1, RAB27A, UNC13D, and STXBP2^10^. Whereas these proteins are directly involved in granule formation, trafficking, or release ^24–27^, KMT2D maintains a transcriptional program required for efficient NK cell release of effector molecules, in part by regulating RAB3D levels. Thus, KMT2D loss of function compromises NK cell secretory machinery through defects in epigenetic regulation.

Comparison of mouse and human RNA-seq data identified a conserved set of KMT2D-dependent genes enriched for exocytosis and vesicle-mediated transport. KMT2D knockout reduced global H3K4me1 abundance and altered H3K4me1 occupancy at regulatory regions associated with this transcriptional program in mature human NK cells. Among these genes, RAB3D functionally links KMT2D-dependent chromatin regulation and NK cell release of effector molecules. KMT2D loss reduced RAB3D transcript and protein levels, and RAB3D loss impaired target-cell killing, degranulation, and extracellular accumulation of GZMB and IFN-γ. The effects on both granule and cytokine release are notable because these pathways use spatially and mechanistically distinct secretory routes in NK cells^5,6,28–31^. RAB3D may therefore represent a shared requirement for these otherwise distinct pathways, consistent with the established functions of RAB3-family proteins in regulated exocytosis in other secretory cell types^21–23,32,33^. However, future work will be necessary to determine the molecular mechanisms by which RAB3D promotes exocytosis or vesicular-mediated transport in NK cells.

The identification of a KMT2D-dependent secretory program in NK cells may also provide a broader framework for understanding the coexistence of neurodevelopmental and immune abnormalities in humans. Although neurons and NK cells perform distinct physiological functions, both require tightly regulated vesicle trafficking and stimulus-dependent exocytosis: neurons release neurotransmitters through synaptic vesicles, whereas NK cells secrete cytokines and cytotoxic granules. RAB3-family GTPases are central regulators of vesicle organization and regulated exocytosis in neuronal and other secretory systems^22,23,33^. Our findings extend these observations by identifying RAB3D as a KMT2D-regulated factor required for efficient human NK-cell degranulation and cytokine release. These observations raise the possibility that KMT2D establishes cell-type-specific secretory competence through epigenetic regulation of RAB3-family proteins or related vesicle-trafficking networks. Disruption of these programs could contribute to pathogenic phenotypes across tissues with high secretory demands, providing a potential mechanistic connection between the neurological, endocrine, and immunological manifestations of KMT2D loss-of-function mutations in individuals with Kabuki Syndrome Type 1^34–38^. Future studies will be necessary to determine whether KMT2D regulates distinct RAB3 paralogs in neurons, immune cells, and other secretory tissues to establish whether defective regulated exocytosis is a shared feature of Kabuki syndrome. More broadly, our study defines chromatin regulation as an upstream determinant of NK cell release of effector molecules and suggests that secretory dysfunction may be an underrecognized consequence of neurodevelopmental disorders caused by mutations in epigenetic regulators.

### Limitations of the study

Although our study identifies KMT2D as a regulator of mature human NK cell effector functions, only two patients have been identified to date with variants in KMT2D that present with an immunodeficiency. Future studies of additional patients will be required to better understand how germline mutations in KMT2D contribute to human NK cell dysfunction through perturbations in development and/or mature NK cell effector functions. Furthermore, although genes associated with effector molecule release and regulated by KMT2D are not changed in the absence of UTX in mouse NK cells, this does not rule out a potential role for KMT2D in regulation of UTX levels and the development of female NK cells in mice and humans. Additionally, while we have shown that RAB3D is required for effector molecule release in human NK cells, it remains possible that RAB3D is one component of a broader KMT2D-dependent secretory program in NK cells rather than its sole downstream effector.

## RESOURCE AVAILABILITY

### Lead Contact

Correspondence and material requests should be directed to Timothy E. O’Sullivan, PhD.

### Materials Availability

This study did not generate any unique reagents.

### Data and Code Availability

All uniquely generated datasets and the corresponding raw files will be deposited Gene Expression Omnibus (GEO) upon publication. UTX knockout RNA-seq and UTX CUT&Tag are published^19^ and accessible as GEO series: GSE185065. All other data are available in the main text or supplemental figures. No custom code was generated for this study.

## ACKNOWLEDGEMENTS

The authors would like to thank the UCLA Center for AIDS Research (CFAR) Virology Core for providing healthy donor peripheral blood samples, the families of NKD patients who donated samples, and the UCLA Broad Stem Cell Research (BSCRC) Sequencing Core for assistance with generating sequencing datasets. T.E.O. is supported by the National Institutes of Health (NIH) (R01AI174519, R01AI186079) and the UCLA Immunology Advisory Committee. K.R.K is supported by the Ruth L. Kirschstein National Research Service Award (AI007323). J.S.O. is supported by the NIH (R01AI120989, R37AI067946). The UCLA CFAR Virology Core is supported by the NIH (5P30AI028697).

## AUTHOR CONTRIBUTIONS

T.E.O. conceived and designed the project, supervised experiments, and wrote the manuscript. K.R.K. performed experiments, analyzed data, generated figures, and wrote the manuscript. G.R.L. analyzed data and generated figures. J.H.J, S.Z., C.G.B, D.H., and L.A.P. performed experiments and analyzed data. J.S.O. provided NKD patient samples and supervised experiments. M.A.S. provided *Kmt2d^fl/f^* mice and supervised experiments. E.H. provided clinical coordination for human samples.

## STAR METHODS

### Mouse colony maintenance

All mouse lines were bred and maintained according to guidelines set forth by the Institutional Animal Care and Use Committee (IACUC) and UCLA Animal Research Committee (ARC). All protocols were approved by IACUC and ARC. Mice used for experiments were between 8-10 weeks of age and sex-matched littermates were used where applicable. C57BL/6J CD45.2^+^ (Jackson) and B6.SJL CD45.1^+^ (Jackson) mice were initially purchased from Jackson Labs and maintained in-house. CD45.1 and CD45.2 mice were crossed to generate CD45.1.2 mice. *Ncr1*^Cre/+^ and *Kmt2d*^fl/fl^ mice were crossed to generate *Ncr1*^Cre/+^*Kmt2d*^fl/fl^ and *Ncr1*^Cre/+^*Kmt2d*^fl/+^ mouse lines.

### Generation of mixed-bone marrow chimeric mice

Mixed bone marrow chimeric mice were generated as previously described^39^. In brief, CD45.1.2 mice were treated 25 mg/kg busulfan intraperitoneally (i.p.) daily for 3 consecutive days. Bone marrow was isolated from WT (CD45.1) or *Ncr1*^Cre/+^*Kmt2d*^fl/+^ (CD45.2), mixed at a 1:1 ratio, and injected into depleted CD45.1.2 mice intravenously (i.v.) via tail-vein. Mice were bled at six weeks post-transfer to evaluate reconstitution efficacy before experimental use.

### Mouse NK cell isolation and enrichment

Mice were euthanized and spleens, livers, and blood were harvested and dissociated as described previously^40^. Single cell suspensions were stained for flow cytometry analysis or used for NK isolation. Briefly, spleens from euthanized mice were removed and dissociated into a single cell suspension in PBS. Homogenized cells were filtered through 100 μM cell strainers, spun down at 1500 rpm for 3 minutes and were resuspended in RoboSep buffer (Stem Cell). NK cells were isolated using negative magnetic selection following the EasySep Mouse NK Cell Isolation Kit (Stem Cell). For *ex vivo* assays, purified NK cells were cultured in complete media (CM) (RPMI 1640, 10% FBS, 2 mM L-glutamine, 2 mM sodium pyruvate, 1% Non-essential amino acids, 1% penicillin/streptomycin, 0.5% sodium bicarbonate, and 55 μM 2-mercaptoethanol) with recombinant mouse IL-15 (50 ng/mL).

### Mouse NK cell effector assays

For plate-bound stimulation assays, purified NK cells were incubated in CM on pre-coated plates for 4 hours in the presence of brefeldin A (BFA) (1:1000), monensin (1:1000), and anti-CD107a antibody (1:400). Plates were prepared using ELISA high-binding plates incubated overnight at 4°C with anti-NK1.1 (clone PK136, 5 μg/mL) or anti-Ly49H (clone 3D10, 5 μg/mL) resuspended in carbonate binding buffer, or with binding buffer alone. NK cells used in cytokine stimulation assays were cultured in CM supplemented with BFA, monensin, recombinant IL-15 (50 ng/mL), and recombinant mouse IL-12 (20 ng/mL) or IL-18 (10 ng/mL) for 4 hours. All assays were performed at 37°C with 5% CO_2._

### Murine Cytomegalovirus Infection

MCMV (Smith strain) was passaged in BALB/C mice three times. On the final passage, MCMV was isolated from salivary glands as previously described^39^. Mice were infected with a sublethal dose, 7.5 x 10^3^ plaque-forming units in 0.5 mL PBS, administered intraperitoneally (i.p.). Mouse survival and body weight were recorded daily. Mice were euthanized if their body weight reached less than 80% of their initial weight (at onset of infection). Immune responses were evaluated at day 1.5 p.i.. at which point mice were euthanized and tissues were harvested as described above.

### Human NK Cell Culture

NKD patient samples were collected in the generation of the NEAR cohort upon approval from the Internal Review Board at the Children’s Hospital of Philadelphia. Patient samples and information were processed and stored according to published criteria^10^. Primary human NK cells were isolated from PBMCs as previously described^19^. Healthy donor PBMCs were obtained from the UCLA Center for AIDS Research (CFAR) Virology Core. NK cells were purified by negative bead selection using the EasySep Human NK Cell Isolation Kit (Stem Cell) and cultured in NK MACS medium (Miltenyi) supplemented with recombinant human IL-2 (200 IU/mL) and IL-15 (20 ng/mL). Cells were expanded for at least 7 days before experimental use.

### Human NK Cell Effector assays

For plate-bound degranulation assays, plates were prepared with anti-NKp30 (clone P30-15, 5 μg/mL) or binding buffer alone. NK cells were incubated in CM with recombinant human IL-2 (200 IU/mL) and IL-15 (20 ng/mL) on pre-coated plates for 12 hours, followed by the addition of BFA, monensin, and anti-CD107a antibody and incubation for an additional 4 hours. For cytokine stimulation assays, NK cells were cultured in CM with IL-2, IL-15 and K562 cells (2.5:1 effector:target ratio), with the addition of recombinant human IL-18 (100 ng/mL) for 16 hours. For tumor-conjugate formation assays a 1:1 ratio of human NK cells and K562-GFP tumor cells were incubated in U-bottom plates for 15 minutes in CM with IL-2 and IL-15. All assays were performed at 37°C with 5% CO_2._

### NK cell Tumor Killing Assays

B2M-deficient MC38 tumor cells were incubated with CellTrace^TM^ Violet (CTV) for 10 minutes, washed with PBS, and resuspended in CM. Purified mouse NK cells were co-cultured with labeled MC38 in CM at an effector to target ratio of 1:2 for 16 hours at 37°C. Wells were washed with PBS, treated with trypsin, and total well contents were collected for flow cytometry. Viable cell counts were based on CTV^+^ gating. Purified human NK cells were cultured in CM with human IL-2, IL-15, and K562-GFP cells at an effector to target ratio of 1:4 for 16 hours at 37°C. Remaining well contents were collected for flow cytometry. Viable cell counts were based on GFP^+^ gating. The quantity of surviving cells was determined via flow cytometry (CTV^+^ or GFP^+^). Specific lysis (%) was calculated as follows: (1 -#tumor cells in co-culture wells /#tumor cells in control well) x 100

^51^Cr-release assays were performed in a 96-well U-bottom plate (Corning) using 1×10^4^ K562 target cells per condition, as previously described^41^. PBMCs (effector cells) were used in serial dilution starting at an effector-to-target cell ratio of 50:1. Cells were co-cultured for 4 hours at 37°C. Supernatants were then transferred into LumaPlates (Revvity) and analyzed using a TopCountXL (PerkinElmer). Specific lysis (%) was calculated as follows: (experimental cpm –spontaneously released cpm)/(total cpm –spontaneously released cpm) x 100.

### Guide RNA design

Guide sequences were designed using published databases. For cRNP editing, guide sequences were selected from previously published genome-wide sgRNA libraries^42^. Cas9 base editing guides were designed using SpliceR (version 1.2.3)^43^. Guide efficacy was evaluated using the EditR tool^44^ to evaluate genomic modifications (*KMT2D*, 15176A>G, *ADAP1, BAIAP2*) or by Western blot to evaluate protein depletion (*RAB3D*).

### CRISPR cRNP editing

cRNP editing was performed as previously described^39,45^. Briefly, sgRNAs (120 pmol, Synthego/Invitrogen) were combined with Alt-R electroporation enhancer (9 pmol, IDT) and water. SpCas9 (20 pmol, qb3 UC Berkley) was diluted in water and combined with the sgRNA solution and allowed to complex for 10 minutes at room temperature. Purified NK cells were spun down at 1500 rpm for 3 minutes, washed one time with PBS, and resuspended in T buffer (Thermo Fisher) at a concentration of 1×10^6^ cells/100 μL T buffer. NK cells were added to complexed cRNP and electroporated using the Neon NxT Transfection System (Thermo Fisher) under the following conditions: 1900 V, 20 ms pulse width, and 1 pulse. Electroporated cells were rested in CM for 60 minutes before centrifugation and replating in either CM with mouse IL-15 (mouse NK cells) or NK MACS with human IL-2 and IL-15 (human NK cells).

### CRISPR Cas9 base editing

Cas9 based editing was performed as previously described^13,19^. Briefly, sgRNAs were mixed with Alt-R electroporation enhancer, water, Protector RNAse Inhibitor (Roche) and mRNA encoding ABE8e cas9 base editor (TriLink Genomics) derived from plasmids provided by the Moriarity laboratory. NK cells were spun down and resuspended in T buffer and added to ABE8e-sgRNA mixes. Cells were electroporated using the Neon NxT system under the following conditions: 1800 V, 10 ms pulse width, and two pulses. Electroporated cells were rested and replated in NK MACS with human IL-2 and IL-15.

### Flow Cytometry

Cells were stained for surface markers/intracellular proteins using fluorophore-conjugated antibodies. Cell surface staining was performed in PBS and cells were stained for 30 minutes at 4°C. Fixation and permeabilization was done with the BD Cytofix/Cytoperm kit for Prf1/PRF1, Gzmb/GZMB, and IFN-γ and eBioscience Foxp3/Transcription Factor kit for UTX and H3K4me1, following manufacturers instructions. Flow cytometry was performed using the Attune NxT Acoustic Focusing cytometer (Thermo Fisher) and data were analyzed with FlowJo v10. Staining was performed using the following antibodies: mouse anti-CD45.1 (1:400, A20), mouse anti-CD45.2 (1:400, 104), mouse anti-CD3 (1:100, 17A2), mouse anti-TCR-β (1:100, H57-597), mouse anti-NK1.1 (1:50, PK136), mouse anti-CD27 (1:200, LG.3A10), mouse anti-CD11b (1:200, M1/70), mouse anti-IFN-γ (1:100, XMG1.2), anti-mouse/human GzmB (1:100, GB11), mouse anti-CD107a (1:100, 1D4B), mouse anti-KLRG1 (1:400, 2F1/KLRG1), mouse anti-Ly49H (1:400, 3D10), human anti-CD3 (1:400, UCHT1), human anti-CD56 (1:200, TULY56), human anti-IFN-γ (1:100, B27), human anti-PRF1 (1:400, B-D48), human anti-CD107a (1:100, H4A3), human anti-CD16 (1:400, CB16), human anti-CD57 (1:100, HNK-1), anti-Mono-Methyl-Histone H3 (Lys4) (1:100, D1A9), anti-UTX (1:100, D3Q1l), goat anti-rabbit H&L (ab6717).

### ELISA

Supernatant from human NK cell effector assays was harvested and stored at -80°C. IFN-γ was detected using an ELISA Max Standard kit (BioLegend) and GZMB were detected using ELISA Max Deluxe kit (BioLegend) following manufacturer’s instructions.

### Western Blots

To extract protein, edited human NK cells were resuspended in Pierce RIPA buffer (Thermo Fisher) with Halt protease inhibitor cocktail (1:100, Thermo Fisher). Protein quantification was done using the Pierce BCA Protein Assay kit (Thermo Fisher). Samples were loaded onto NuPage Novex 4-12% Bis-Tris Protein Gels, electrophoresed, transferred to PVDF membranes, and blocked overnight with 5% milk resuspended in 1x TBS with 0.1% Tween-20 (TBST) at 4°C. Membranes were washed in TBST and immunoblotted with rabbit anti-RAB3D (1:1000, EPR8106) or rabbit anti-β-actin (1:2000, 13E5) diluted in 5% milk-TBST for 1 hour at room temperature (RT). Secondary staining was performed with goat anti-rabbit horseradish peroxidase (1:10,000, CST4970) diluted in 5% milk-TBST for one hour at RT. Proteins were visualized using the SuperSignal West Pico PLUS ECL kit (Thermo Fisher) and visualized using the Azure Biosystems c280 imager. Western blots were quantified using ImageJ.

### RNA-seq library construction

Total RNA was isolated from mouse or human NK cells using the Zymo Quick-RNA MicroPrep kit (R1051) according to manufacturer instructions. RNA concentration and quality were assessed using a NanoDrop spectrophotometer and Qubit fluorometer (Thermo Fisher Scientific). Mouse libraries were prepared as previously described^46^. Sequencing was performed at the Broad Stem Cell Research Center (BSRSC) Sequencing Core at UCLA and run on a NovaSeq X platform (Illumina) generating paired-end reads of 2×100 bp. The targeted sequencing depth was 100 million reads per sample. Human library sequencing was performed using the Plasmidsaurus RNA-Seq pipeline from frozen RNA samples at a minimum concentration of 10 ng/μL. Libraries were run on a NovaSeq X Platform (Illumina) using a 3’ end counting approach, with an aim of returning 10 million deduplicated reads from 20 million raw reads.

### RNA-seq analysis

Raw paired-end (mouse) or single-end (human) reads from the Illumina NovaSeq X platform were processed to generate gene expression counts. Initial quality control of the raw FASTQ files was performed using FastQC (version 0.12.0). Adapter trimming and low-quality base removal were conducted using Trimmomatic (version 0.39)^47^. Specifically, paired-end reads were trimmed to remove Illumina Nextera adapters using the ILLUMINACLIP parameter with the adapter sequence file “NexteraPE-PE.fa” and the following settings: 2:30:10. Additionally, a sliding window trimming approach (SLIDINGWINDOW:4:20) was applied to remove low-quality bases from the ends of reads, and reads shorter than 40 bp were discarded (MINLEN:40). Leading and trailing low-quality bases (quality score < 3) were also removed (LEADING:3 TRAILING:3). The trimmed paired-end reads were then aligned to either the GRCh38 reference genome for human or GRCm39 reference genome for mouse using STAR aligner (version 2.7.11)^48^. The STAR alignment was performed with the parameter --quantMode GeneCounts to obtain ‘.ReadsPerGene.tab’ gene counts files for each sample. Following read alignment and gene counts calculation, differential gene expression analysis was performed using DESeq2 (version 1.42.1)^49^. Human samples were analyzed in a paired manner using the formula ∼condition+patientID in the DESeqDataSetFromMatrix() function to account for inter-patient variability, while mouse samples were analyzed in a non-paired manner using the formula ∼condition. Genes with a padj < 0.05 were considered statistically significant, and a minimum |log2(fold change)| of 0.5 (human) or 0.7 (mouse) was applied to identify genes with biologically relevant changes in expression. Cross-species gene comparisons were made using biomaRt^50^, genes were filtered by padj and log2 fold change, and considered conserved if log2 fold change was shared directionally. The results of the differential gene expression analysis were visualized using volcano plots generated with ggplot2 (version 3.5.1)^51^. Functional enrichment analysis was performed using g:Profiler^52^. All R packages were run using R version 4.6.1 in R Studio.

### CUT&Tag Library Preparation

CUT&Tag library preparation for anti-H3K4me1 and anti-H3K27ac was performed as previously described^19,46^. Nuclei were isolated from edited human NK cells with cold nuclear extraction buffer (20 mM HEPES pH 7.9, 10 mM KCl, 0.1% Triton X-100, 20% glycerol, 0.5 mM spermidine diluted in Ix protease inhibitor cocktail -Roche) and incubated with activated concanavalin A-coated magnetic beads (Polysciences, 86057-3) for 3 minutes at RT. Antibodies for H3K4me1 (Mono-Methyl-Histone H3 Lys4; D1A9, Cell Signaling Technology), H3K27ac (Acetyl-Histone H3 Lys27; D5E4, Cell Signaling Technology), or IgG isotype control (3900S, Cell Signaling Technology) were diluted in antibody buffer (20 mM HEPES pH 7.5, 150 mM NaCl, 0.5 mM spermidine, 1x protease inhibitor cocktail, 0.05% digitonin, 2mM EDTA, 0.1% BSA) and added to isolated nuclei. Samples were incubated overnight at 4°C on a rotator. The next day, samples were placed on a magnetic tube holder and supernatants were discarded. Secondary antibody (guinea pig anti-rabbit IgG, NBP172763, Fisher Scientific) was diluted 1:100 in Dig-Wash (20 mM HEPES pH 7.5, 150 mM NaCl, 0.5 mM spermidine, 1x protease inhibitor cocktail, 0.05% digitonin) and incubated for an hour at RT. Nuclei were washed four times in Dig-Wash and incubated with pAG-Tn5 adaptor complex (EpiCypher) in Dig-300 buffer (20 mM HEPES pH 7.5, 300 mM NaCl, 0.5 mM spermidine, 1x protease inhibitor cocktail) for 1 hour at RT. Tagmentation was stopped by addition of Dig-300 buffer with 10 μL 1 M MgCl, 7.5 μL 0.5 M EDTA, 2.5 μL 10% SDS and 5 μL 10 mg/mL proteinase K. Samples were incubated for 1 hour at 55°C. DNA was extracted via phenol:chloroform:isoamyl alcohol separation. DNA was barcoded using Illumina i7 primers and amplified using the following conditions: PCR mix containing 25 μL NEBNext 2x mix, 2μL each of forward and reverse 10 μM primers and 21 μL extracted DNA. Amplification conditions: 58°C for 5 minutes, 72°C for 5 minutes, 98°C for 45 seconds, 14x 98°C for 15 seconds followed by 63°C for 10 seconds, 72°C for 1 minute. Amplified libraries were purified using KAPA pure SPRI beads (Roche) and DNA was eluted in TE buffer. All libraries were mixed at equimolar proportions. Sequencing was performed at the UCLA BSRSC sequencing core and run on a NovaSeq X Plus sequencer (Illumina), generating 2×100 bp paired-end reads with a target depth of 100 million reads per sample.

### CUT&Tag analysis

Raw paired-end sequencing reads were subjected to quality control using FastQC (version 0.12.0). Adapter trimming and removal of low-quality bases were performed using Trimmomatic (version 0.39)^47^ as described above. Trimmed reads were aligned to the human genome (hg38), downloaded from UCSC and indexed using Bowtie2-build, using Bowtie2 (version 2.5.4)^53^ with the following parameters: ‘*--local --very-sensitive --no-mixed --no-discordant --phred33 -I 10 -X 700*’. Alignment files were converted to sorted BAM format using SAMtools (version 1.21)^54^. Reads mapping to mitochondrial DNA and blacklisted regions were removed using SAMtools. BigWig files for visualization were generated using deepTools (version 3.5.6)^55^ ‘*bamCoverage*’ with default parameters. Significant enrichment regions (peaks) were identified using SEACR (version 1.3)^56^ with the following parameters: ‘*0.01 norm stringent*’. The non-treated control was used to inform the threshold for significant enrichment using the default normalization (’norm’) and stringent peak calling (’stringent’) settings with a q-value cutoff of 0.01. Peak annotation to the nearest gene was performed using the ‘annotatePeaks.pl’ function from HOMER (version 5)^57^ with the corresponding genome assembly (’*-genome hg38*’ downloaded via HOMER) and default parameters. Unique gene names associated with significantly bound regions (by either H3K4me1 or H3K27ac) were extracted for downstream analysis.

### Statistical Analysis

Statistical comparisons between groups were performed using two-tailed Student’s T tests with Welch’s correction, Wilcoxon matched-pairs signed rank test, two-way ANOVA with multiple comparisons, performed with GraphPad Prism Software (v11.0.2) unless otherwise noted in the figure legends. All graphs display datapoints representing individual donors or mice, with connected points originating from the same donor. All error bars represent the standard error of the mean (S.E.M.). The p-value significance threshold was set to p < 0.05 for all analyses. Researchers were unblinded to treatment groups when performing experiments and assessing data. Preliminary and previously published data informed sample size selection to ensure adequate statistical power.

