## Supplemental Figures for "The Histone Methyltransferase KMT2D promotes Natural Killer cell effector molecule release"

### SUPPLEMENTAL INFORMATION

**Figure S1.** Profiling KMT2D-deficiency in human NK cells (related to Figure 1).

**Figure S2.** *Kmt2d* represses mature mouse NK cell development (related to Figure 2).

**Figure S3.** Epigenetic regulation of human NK cells by KMT2D (related to Figure 3).

**Figure S4.** KMT2D and RAB3D do not control the intracellular levels of effector molecules (related to Figure 4).

### Supplemental Figure 1

**A**

| Patient ID | Coding Exon | Nucleotide Variant | Amino Acid Variant | Zygosity | PLI/AF[1] | CADD[2] |
|---|---|---|---|---|---|---|
| Patient 1 | exon 31 | c.C6752T | p.S2251L | Heterozygous | 1/1.64e-3 | 27.8 |
| Patient 2 | exon 39 | c.11217_11222del | p.Gln3744_Gln3745del | Heterozygous | 1/2.23e-5 | n/a |

[1]Phred-scaled Likelihood/Allele Frequency
[2]Combined Annotation Dependent Deletion

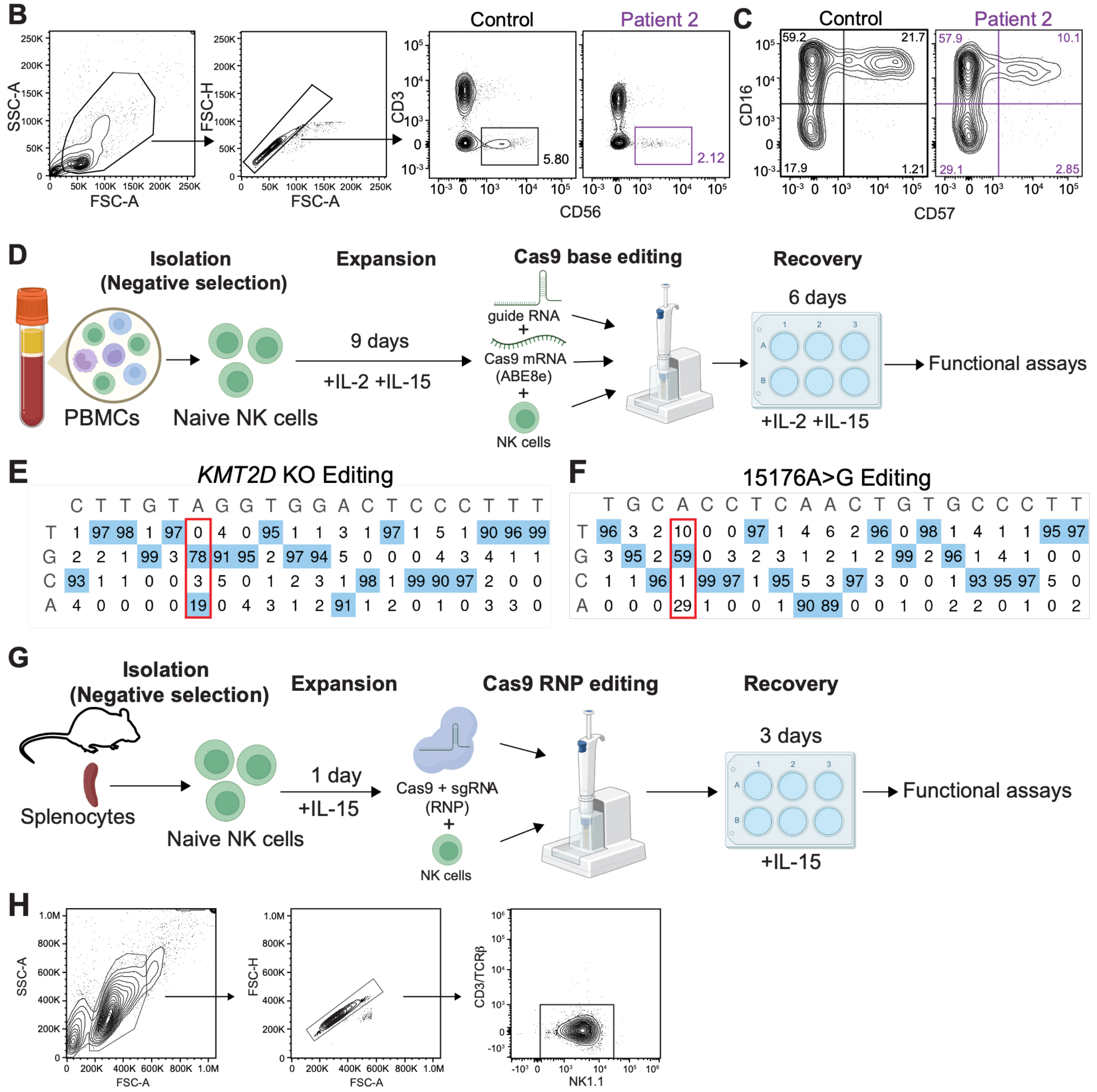


**Supplemental Figure 1. Profiling KMT2D-deficiency in human NK cells (related to Figure 1).** (**A**) Summary table of genetic features from patients with a mutation in *KMT2D*, including nucleic acid and amino acid annotations, zygosity, and Phred-scaled Likelihood (PLI), allele frequency (AF), and Combined Annotation Dependent Deletion (CADD) scores. (**B**) Flow cytometry gating strategy for human NK cells. (**C**) Representative plots of CD16 and CD57 staining in $CD56^+$ PBMCs from Control and NKD Patient 2 samples. (**D**) Schema of Cas9 base editing workflow for human NK cells. (**E,F**) Per position nucleotide frequency from Sanger sequencing of *KMT2D* KO (**E**) or 15176A>G (**F**) CRISPR edits, estimated with EditR[44] with boxes indicating target sites. (**G**) Schema of Cas9 RNP editing workflow for mouse NK cells. (**H**) Flow cytometry gating strategy for mouse NK cells.

### Supplemental Figure 2

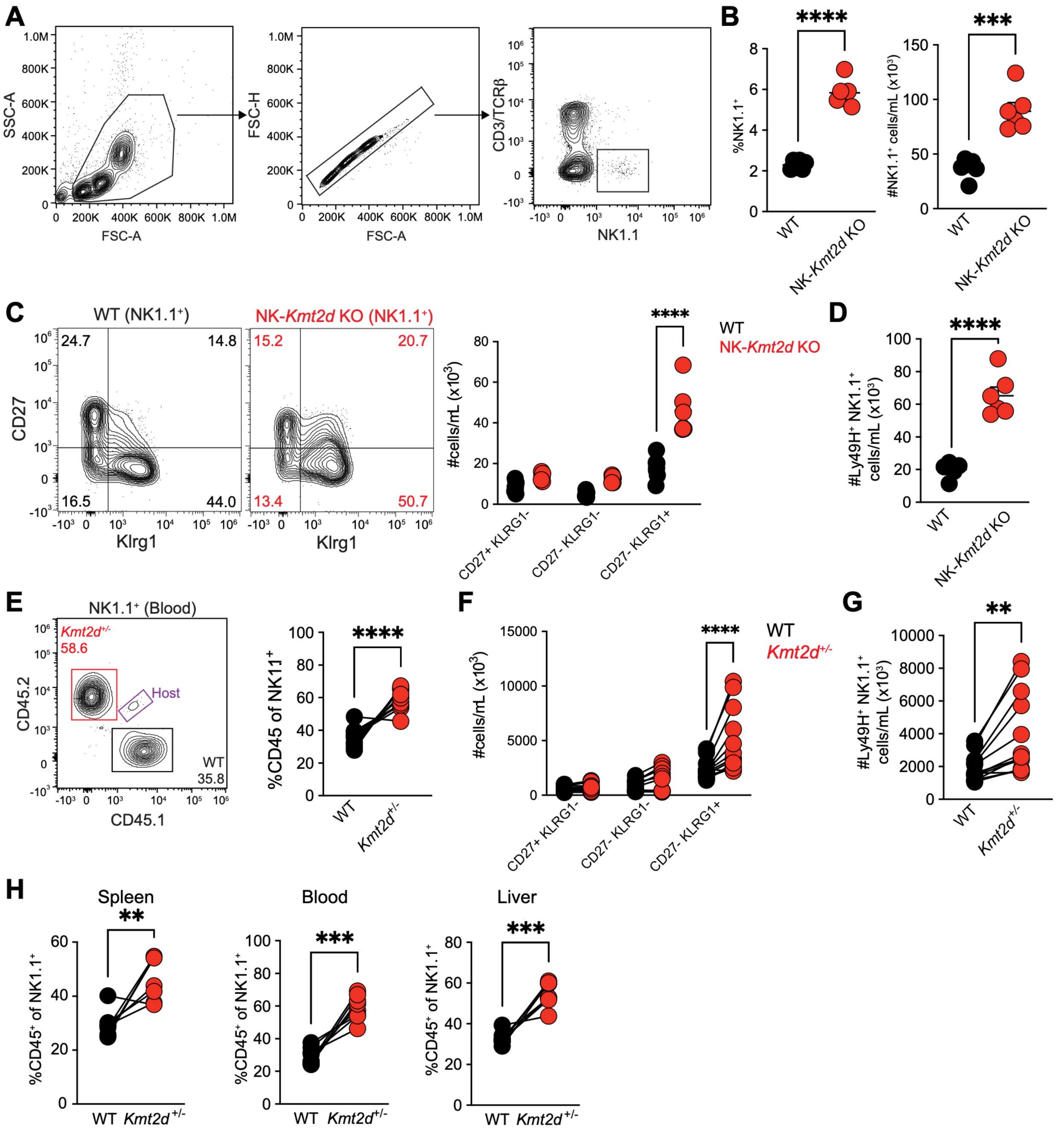


**Supplemental Figure 2. *Kmt2d* represses mature mouse NK cell development (related to Figure 2).** (**A**) Gating strategy for mouse NK cells. (**B**) Percent (left) and number (right) of NK1.1$^+$ cells in blood of uninfected NK-*Kmt2d* KO versus WT mice. (**C**) Representative plot (left) and quantification (right) of CD27 and Klrg1 from the blood of uninfected NK-*Kmt2d* KO versus WT mice. (**D)** Number of Ly49H$^+$ NK1.1$^+$ cells in the blood of uninfected NK-*Kmt2d* KO versus WT mice. (**E**) Representative plot of CD45.1 and CD45.2 staining (left) and proportion (right) of WT versus *Kmt2d*$^{+/-}$ NK cells in the blood of uninfected mBMCs. (**F**) Quantification of CD27 and Klrg1 NK subsets from *Kmt2d*$^{+/-}$ versus WT cells in the blood of uninfected mBMCs. (**G**) Number of Ly49H$^+$ NK1.1$^+$ cells derived from *Kmt2d*$^{+/-}$ versus WT cells in the blood of uninfected mBMCs. (**H**) Quantification of the proportion of WT versus *Kmt2d*$^{+/-}$ NK cells in the spleen (left), blood (middle), and liver (right) of mBMCs at day 1.5 post-MCMV infection. Data represent (**B-D**) mean ± SEM or (**E-H**) paired individuals, and at least two independent experiments with (**B-D**) n=6, (**E-G**) n=13, or (**H**) n=5 mice. *p<0.05, **<0.01, ***<0.001, ****<0.0001 by (**B,D**) Student's T-test (**C**) two-way ANOVA, (**E,G,H)** Wilcoxon matched-pairs sign rank test, or (**F**) paired two-way ANOVA.

### Supplemental Figure 3

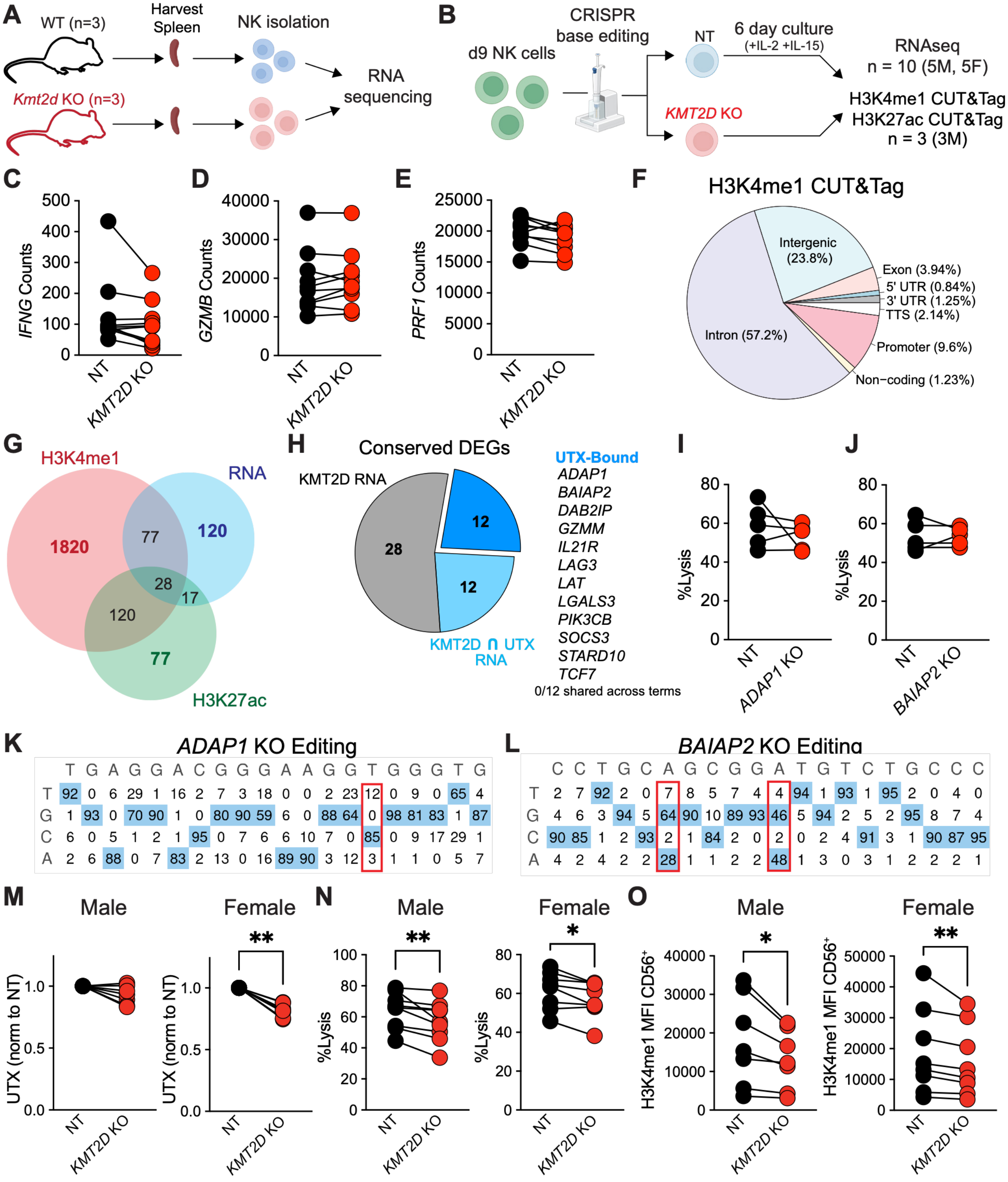

**Supplemental Figure 3. Epigenetic regulation of human NK cells by KMT2D (related to Figure 3).** (**A,B**) Experimental strategy for generating mouse RNA-seq (**A**) and human RNA-seq, H3K4me1 CUT&Tag, and H3K27ac CUT&Tag (**B**) datasets. (**C-E**) RNA-seq transcript counts in *KMT2D* KO versus NT human NK cells of *IFNG* (**C**), *GZMB* (**D**), and *PRF1* (**E**). (**F**) Pie chart showing the proportion of *KMT2D*-dependent H3K4me1 binding within different genomic regions in human NK cells. (**G**) Venn diagram showing the overlap between RNA-seq DEGs and differentially bound peaks in H3K4me1 and H3K27ac CUT&Tag in *KMT2D* KO versus NT human NK cells. (**H**) Chart showing the number of conserved DEGs dependent on KMT2D, UTX, and bound by UTX (left) and the list of UTX-bound genes (right). (**I,J**) Specific lysis of K562-GFP tumor cells after 16hr co-culture with *ADAP1* KO (**I**) or *BAIAP2* KO (**J**) versus NT controls at a 1:4 E:T in the presence of IL-2 and IL-15. (**K,L**) Per position nucleotide frequency from Sanger sequencing of *ADAP1* KO (**K**) or *BAIAP2* KO (**L**) CRISPR edits, estimated with EditR[44] with boxes indicating target sites. (**M**) UTX MFI (normalized to NT) in $CD56^+$ NT versus *KMT2D* KO human NK cells in male (left) and female (right) donors. (**N**) Percent specific lysis of K562-GFP tumor cells after 16hr co-culture with *KMT2D* KO versus NT NK cells from male (left) and female (right) donors at a 1:4 E:T in the presence of IL-2 and IL-15. (**O**) H3K4me1 MFI in $CD56^+$ NT versus *KMT2D* KO human NK cells in male (left) and female (right) donors. Data represent paired human donors with (**C-E**) n=10, (**I,J**) n=5, or (**M-O**) n=7-9 donors and at least two independent experiments. *p<0.05, **<0.01 by Wilcoxon matched-pairs sign rank test.

### Supplemental Figure 4

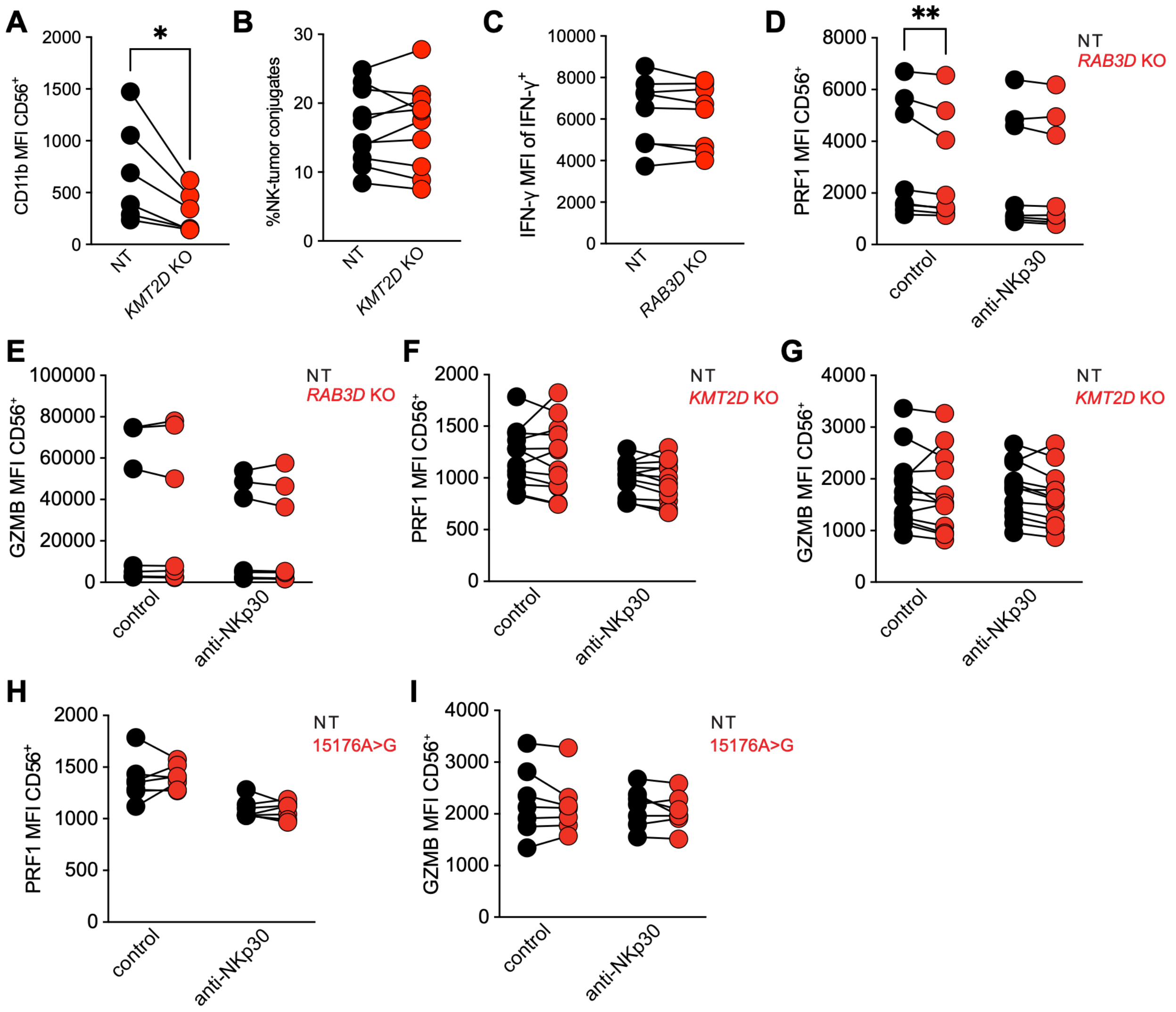


**Supplemental Figure 4. KMT2D and RAB3D do not control the intracellular levels of effector molecules (related to Figure 4).** (**A**) MFI of CD11b in *KMT2D* KO versus NT human NK cells. (**B**) Percentage of *KMT2D* KO versus NT NK cells conjugated to K562-GFP tumor cells after 15 minutes of co-culture in the presence of IL-2 and IL-15. (**C)** MFI of IFN-γ in IFN-$γ^+$ $CD56^+$ *RAB3D* KO versus NT human NK cells after 16hr co-culture with K562-GFP in the presence of IL-2, IL-15, and IL-18. (**D-I**) MFI of PRF1 (**D, F, H**) or GZMB (**E,G,I**) in $CD56^+$ *RAB3D* KO (**D,E**), *KMT2D* KO (**F,G**), or 15176A>G (**H,I**) versus NT human NK cells after 16hr stimulation with plate-bound anti-NKp30 or control conditions in the presence of IL-2, IL-15, and 4 hours of anti-CD107a, BFA, and Monensin. Data represent paired individuals and at least two independent experiments with (**A**) n=6, (**B**) n=10, (**C-E**) n=8, (**F,G**) n=12, or (**H,I**) n=7 by paired 2-way ANOVA. *p<0.05, **<0.01 by (**A-C**) Wilcoxon matched-pairs sign rank test or (**D-I**) paired 2-way ANOVA.
